# Enhancing the interoperability of glycan data flow between ChEBI, PubChem, and GlyGen

**DOI:** 10.1101/2021.06.17.448729

**Authors:** Rahi Navelkar, Gareth Owen, Venkatesh Mutherkrishnan, Paul Thiessen, Tiejun Cheng, Evan Bolerlton, Nathan Edwards, Michael Tiemeyer, Matthew P Campbell, Maria Martin, Jeet Vora, Robel Kahsay, Raja Mazumder

## Abstract

Glycans play a vital role in health, disease, bioenergy, biomaterials, and biotherapeutics. As a result, there is keen interest to identify and increase glycan data in bioinformatics databases like ChEBI and PubChem, and connecting them to resources at the EMBL-EBI and NCBI to facilitate access to important annotations at a global level. GlyTouCan is a comprehensive archival database that contains glycans obtained primarily through batch upload from glycan repositories, glycoprotein databases, and individual laboratories. In many instances, the glycan structures deposited in GlyTouCan may not be fully defined or have supporting experimental evidence and citations. Databases like ChEBI and PubChem were designed to accommodate complete atomistic structures with well-defined chemical linkages. As a result, they cannot easily accommodate the structural ambiguity inherent in glycan databases. Consequently, there is a need to improve the organization of glycan data coherently to enhance connectivity across the major NCBI, EMBL-EBI, and glycoscience databases.

This paper outlines a workflow developed in collaboration between GlyGen, ChEBI, and PubChem to improve the visibility and connectivity of glycan data across these resources. GlyGen hosts a subset of glycans (~29,000) from the GlyTouCan database and has submitted valuable glycan annotations to the PubChem database and integrated over 10,500 (including ambiguously defined) glycans into the ChEBI database. The integrated glycans were prioritized based on links to PubChem and connectivity to glycoprotein data. The pipeline provides a blueprint for how glycan data can be harmonized between different resources. The current PubChem, ChEBI, and GlyTouCan mappings can be downloaded from GlyGen (https://data.glygen.org).

## INTRODUCTION

Glycosylation is one of the most abundant, complex and diverse post-translational phenomena. The process by which aglycones mainly proteins or lipids are modified by glycans to form glycojugate is ubiquitous and essential to all life forms. Significant research has taken place to study the synthesis, structure, function, and metabolism of glycans for its application in numerous diverse fields (O’Neill et al. 2015; Penades et al. 2015; Reily et al. 2019; Seeberger and Cummings 2015).

### Role of glycans

The main driver for the glycobiology research has been human health and disease. This is primarily due involvement of the process in many important biological functions like transcription, translation, cell division, cellular differentiation, tissue development, cell adhesion, cell-cell interactions, protein folding and stability, inflammatory responses and many others (Varki 2017; Varki and Gagneux 2015). Altered glycosylation can be seen in different diseases like cancer, autoimmune disorders (Rheumatoid arthritis, IBS, etc.), and others. This has encouraged researchers to identify and study the differential glycosylation (i.e. alterations to the sialyation, fucosylation, branching, structure, etc.) for development of diagnostic and prognostic biomarkers. Examples include analyzing core fucosylation and protein concentration of alpha fetoprotein L3 as biomarker for hepatocellular carcinoma (Kirwan et al. 2015; Shiraki et al. 1995) or profiling altered glycoforms on MUC16 and MUC1 for ovarian cancer (Chen et al. 2013). Glycans and glycoproteins are also involved in host-pathogen interaction and have many important roles including entry, mediation and proliferation of pathogens into host cells, glycan shielding by pathogens against host’s immune response, utilization of glycoproteins by host as barriers against pathogens as well as triggering an immune response in host upon pathogen recognition. (Crispin et al. 2018; Li et al. 2017; Lin et al. 2020; Miller et al. 2008; Walls et al. 2016; Wang et al. 2004). It has been understood that the viral proteins are extensively glycosylated such as spike glycoprotein for SARS-CoV2 (Watanabe et al. 2020), envelop glycoprotein gp120 in HIV-1 (Panico et al. 2016), which play a vital role in replication and pathogenesis. e.g. mutations in glycosylation’s sites of the NS1 protein impacts the pathogenicity of Dengue 2 virus (DENV-2) (Crabtree et al. 2005). As a result, the involvement of glycosylation is extensively studied to potentially develop vaccine targets and therapies e.g. utilization of Heparin sulfate to inhibit HSV-1 entry and infection (Copeland et al. 2008; Sharthiya et al. 2017).

Applications of glycans is not limited to the biomedical field. For several years glycans have been used in construction and paper making industry, food and beverages, transportation, and textile industry and extensive research is implemented to understand the utility in several fields such as renewable energy production, developing nanomaterials, enhanced polymers as well as generating high-value chemicals (National Research Council 2021; O’Neill et al. 2015). Due to the global climate crisis researchers are interested in finding sustainable and economic alternatives to petroleum-based energy production. One such source is utilizing plant cell walls consisting of diverse complex polysaccharides such as hemicellulose, cellulose and pectin as lignocellulosic biomass for energy production. As a result, several crops are investigated for production of bioenergy which include switchgrass (Lemus et al. 2014; Schmer et al. 2008), sudangrass (Acevedo et al. 2019), eucalyptus (Bonifacino et al. 2021), miscanthus (da Costa et al. 2017), poplar and willow (Clifton-Brown et al. 2019). Additionally, the byproducts developed during the processing of biomass such as xylose or glucose are also being studied for their role in producing functional chemical precursors including ethanol, xylitol, hydroxymethlyfurfural, lactic acid (O’Neill et al. 2015). Another application includes utilizing plant glycans to develop polymers with enhanced properties (Hansen and Plackett 2008; Hartman et al. 2006). Examples include modifying cellulose to generate cellophane, rayon and other derivatives like cellulose butyrate, cellulose acetate (National Research Council 2012; O’Neill et al. 2015). Due to high tensile strength and stiffness provided by the structure of cellulose, it is considered for development of lignocellulose based nanomaterials (referred as CNs) for a wide variety of industry applications including drug delivery systems, cardiovascular implants, different films (such as barrier, transparent and antimicrobial), nanofillers, binders, etc. (Daus and Heinze 2010; Figueiredo et al. 2017; Sai Prasanna and Mitra 2020; Wijaya et al. 2021).

#### Significance of glycoinformatics

Due to the immense diversity of structures generated through process as well as the microheterogeneity it is difficult to link structures with essential functions (Cummings and Pierce 2014; Varki and Gagneux 2015). Thus, it is evident that in-depth knowledge of the structure, function of the glycans as well as the associated conjugates, enzymes and pathways is vital for utilization of glycans and glycoproteins in development of biomarkers, vaccine design, and therapeutics as well as in renewable energy production, biofuels, and nanomaterials at a global and commercial level. The influx of knowledge available through literature needs to be supplemented with biocuration, standardization and translation of raw data as value-added knowledge into open-access bioinformatics databases to facilitate further research. However, a problem facing the bioinformatics community is the availability of published fully-determined glycan structures (where there is complete information of the arrangement of the monosaccharides, linkages/anomeric configuration and glycosidic bonds). Even though it is expected that the glycan analysis provides topology information, it is difficult to transfer such data from literature to bioinformatics databases without a curator’s effort to evaluate composition and other supplementary information to generate fully determined structures. Thus, the knowledge transfer from literature to bioinformatics resources is lengthened significantly. Among the available glycan and glycoconjugate database resources available, the resources focused in the paper are briefly described below.

### GlyTouCan

GlyTouCan (Tiemeyer et al. 2017) (https://glytoucan.org/) designed to serve a function analogous to what GenBank serves for nucleotide sequences provides a repository of uncurated glycans which are fully-defined or with structural ambiguity at multiple levels which can be described by the Glycan naming and subsumption ontology or GNOme ((http://obofoundry.org/ontology/gno, http://gnome.glyomics.org) (Supplementary Figure S1). Depositions made in GlyTouCan may also include monosaccharide compositions without any structural context or structural representations without complete description of the linkages between their monosaccharide building blocks. In addition to hosting over 120,000 glycans (https://glytoucan.org/Structures) and assigning accessions for individual glycans (which are adopted as primary identifiers by glycoprotein-centric databases such as GlyGen (York et al. 2020), GlyConnect (Alocci et al. 2019) and UniCarbKB (Campbell et al. 2014)), GlyTouCan also provides cross-references to contributing databases and includes other annotations such as digital sequences (such as GlycoCT (Herget et al. 2008), WURCS (Matsubara et al. 2017)), literature evidence, and Symbol Nomenclature for Glycans (SNFG) images (Neelamegham et al. 2019; Varki et al. 2015). GlyTouCan provides text or graphic tools, APIs to search or register the glycans as well as SPARQL endpoint (http://code.glytoucan.org/rdf-ontology/sparql-queries/) to download data from the database.

### ChEBI

ChEBI (Degtyarenko et al. 2008) (https://www.ebi.ac.uk/chebi/) is a chemical database of small molecules hosting over 50,000 annotated entries (https://www.ebi.ac.uk/chebi/statisticsForward.do). ChEBI provides unique identifiers for chemical entities (including glycans), host text-string representation (such as SMILES, InChI, InChI key, etc.), 2-D structure of entities, and associated literature evidence. In addition to providing unique IDs for chemical entities, ChEBI links them to other important databases including reaction databases like Rhea (Lombardot et al. 2019) and pathway databases like Reactome (Jassal et al. 2020) that show the individual entities that take part in any given reaction (i.e. the reactants and products). Furthermore, ChEBI also provides ontology (http://www.obofoundry.org/ontology/chebi.html) for chemical entities where groups, classes, child-parent relationships are provided and used by a number of research groups including Gene Ontology (Ashburner et al. 2000; Gene Ontology Consortium 2021) and model species groups. ChEBI provides simple and advances tools to search, view and download individual entries in SDF formats and provides FTP, Oracle and SQL dumps for bulk data download (https://www.ebi.ac.uk/chebi/downloadsForward.do).

### PubChem

PubChem (Kim et al. 2021)(https://pubchem.ncbi.nlm.nih.gov/) is an open chemical database of (primarily) small and large molecules, hosting over 109,000,000 compounds (https://pubchemdocs.ncbi.nlm.nih.gov/statistics) sourced from over 700 resources (https://pubchem.ncbi.nlm.nih.gov/sources). The sourced entities are referred as “substances” and are associated with unique Substance (SIDs) which are then standardized to unique “compound” identifiers (or CIDs). PubChem provides 2-D, 3-D structures, SNFG (SVG) images for glycans, text-string representation (such as IUPAC-Condensed, LINUCS, WURCS, SMILES, InCHI, etc), chemical and physical properties, related/similar compounds, biologic descriptions, synonyms, literature evidence and links to contributing resource(s) for the hosted compounds. Additionally, PubChem provides upload mechanisms where different groups or organizations can submit and update data systematically. Moreover, PubChem provides download options for individual or bulk download via FTP and other programmatic access services (https://pubchemdocs.ncbi.nlm.nih.gov/downloads).

### GlyGen

GlyGen (https://www.glygen.org/) is a glycoinformatics resource with extensive glycoprotein and glycan data integrated from various databases. GlyGen data is primarily divided into protein/glycoprotein and glycan domains where dedicated entry pages are provided (e.g. https://www.glygen.org/protein/P14210-1 and https://www.glygen.org/glycan/G17689DH) for each protein and glycan. To facilitate interoperability, GlyGen uses accessions from UniProtKB (The UniProt Consortium 2021) and GlyTouCan as primary accessions to identify proteins and glycans respectively. Additionally, several cross-references to other protein databases (like RefSeq, PDB, etc), glycoprotein databases (like GlyConnect, UniCarbKB, etc.) and other glycan or chemical databases (such as PubChem, MatrixDB (Clerc et al. 2019), Glycosciences.DB (Bohm et al. 2019), KEGG glycans (Hashimoto et al. 2006), etc.) are provided. Additionally, for glycans unique digital representation (GlycoCT, WURCS, SMILES, IUPAC, GLYCAM IUPAC, etc.), harmonized symbolic nomenclature representations SNFG, and important annotations such as glycan classification (N-linked, O-linked, high mannose, core 1, etc.), glycan motifs, glycosylation annotations (site-specific or global) on proteins (e.g. https://glygen.org/protein/P14210-1#Glycosylation), relevant supporting publications, as well as protein annotations related to disease, expression, mutation, etc. are provided. GlyGen also utilizes Gene ontology, Protein ontology (https://proconsortium.org/) and GNOme. GlyGen provides simple, advanced tool to search and perform complex queries across multiple domains. GlyGen data can be downloaded through GlyGen DATA (https://data.glygen.org/) or via API (https://api.glygen.org/) or SPARQL (https://sparql.glygen.org/) services.

Additionally, GlyGen also provides comprehensive documentation of integration process of individual datasets at GlyGen DATA portal via BioCompute objects (Patel et al. 2021; Simonyan et al. 2017). For example https://data.glygen.org/GLY_000001/v-1.8.25/bco.

It is important to enhance the connectivity across GlyTouCan, ChEBI and PubChem databases by promoting interoperability of glycan data available at GlyGen, provided by its project partners. This will allow GlyGen to connect glycans to glycoproteins, reaction and pathway annotations and ontologies provided by GNOme and ChEBI. This will allow users to navigate across multiple resources and via one portal. Previously, UniCarbKB demonstrated the importance of connecting glyco-centric databases with PubChem with submission of ~1,700 GlycoSuiteDB entries.

## RESULTS

The primary goal of this collaboration is to connect GlyGen, PubChem, ChEBI and GlyTouCan databases at a glycan level and connect them further to reaction and pathways information or resources. The resulting interoperability will allow users to navigate the same glycan across different resources and access information unique to each database through one domain i.e. GlyGen.

### Expanding glycan data coverage in ChEBI

Utilizing the data-flow and collaboration between the databases, over 10,400 glycans were registered or mapped semi-automatically including 10,121 fully-determined and 25 high-value (i.e. reported on large number of glycoproteins, see Supplementary Table SI) ambiguously defined glycans. The integrated glycans are annotated with structure data (for glycans with a PubChem CID cross-reference), formula, charge, mass values, InChI, InChIKey and SMILES. Additionally, new ChEBI names (while retaining the original name as a synonym), more precise ontology terms (such as “amino tetrasaccharide” or acetamides”) are added to generate 3-star curated status entries. The corresponding GlyGen and GlyTouCan cross-references are added under “Manual X-ref” whereas PubChem IDs are listed under “Automatic X-ref” sections of the individual ChEBI entry pages respectively (e.g. https://www.ebi.ac.uk/chebi/searchId.do?chebiId=70967). Among the integrated glycans in ChEBI, ambiguously defined glycans (not previously present in ChEBI) are annotated with ontology, synonyms (in this case WURCS digital sequence) and cross-references to GlyTouCan and GlyGen databases. However, they defer from other entries such that 1) they are without 2-D structure images due to high number of possible linkages and 2) the ChEBI names are generated using a specific format which includes prefix of the source database followed by accession provided by the database (e.g. GlyTouCan G06110VR) e.g. https://www.ebi.ac.uk/chebi/searchId.do?chebiId=156559. List of ambigously defined glycans registered in ChEBI and their corresponding ChEBI IDs can be found in Supplementary Table SI. Additionally, new ontology classes adapting Glycan Naming ontology (or GNOme) were created within ChEBI ontology to classify the amigously defined glycans. Classes such as base-composition (https://www.ebi.ac.uk/chebi/searchId.do?chebiId=CHEBI:167481), composition (https://www.ebi.ac.uk/chebi/searchId.do?chebiId=CHEBI:167502) and topology (https://www.ebi.ac.uk/chebi/searchId.do?chebiId=CHEBI:167503) were created and linked to the corresponding instances in GNOme.

Moreover, as ChEBI routinely submits entries to the PubChem database, any new glycans (expect ambiguously defined glycans) added in ChEBI through this collaboration were also added to the PubChem database. This added either new glycans to PubChem or added a ChEBI cross-reference to glycans already present in PubChem. Additionally, it was observed during the integration process that certain structures (e.g. beta-D-Glucosamine; CID441477) in PubChem were mapped to both its monomeric (e.g. CHEBI:28393 beta-D-Glucosamine) and polymeric (e.g. Chitosan; CHEBI:16261) versions in ChEBI. This was because ChEBI represents polymeric structures where the repeating part of the monomeric structure was represented in square brackets with an “n” multiplier outside of the brackets was ignored during PubChem’s processing. Such mappings were corrected by GlyGen by matching the InChI strings of the entries. The resolved mappings are currently available through GlyGen (e.g. https://glygen.org/glycan/G19577LJ).

This collaboration significantly increased the number of glycans (from ~800 in Feb 2020 to ~10,400) in ChEBI as well as increased the number of PubChem CIDs with CHEBI cross-reference (from ~3729 in April 2020 to ~9409). The integrated glycans can be accessed via GlyGen DATA portal (https://data.glygen.org/GLY_000296) or individual pages (e.g. https://glygen.org/glycan/G78059CC#Cross-References) or ChEBI FTP (ftp://ftp.ebi.ac.uk/pub/databases/chebi/Flat_file_tab_delimited/database_accession.tsv or CHEBI entry pages (https://www.ebi.ac.uk/chebi/searchId.do?chebiId=70967) or PubChem FTP (https://ftp.ncbi.nlm.nih.gov/pubchem/Compound/Extras/CID-Synonym-filtered.gz) or individual entry pages (e.g. https://pubchem.ncbi.nlm.nih.gov/compound/70679232)

### Submitting glycan annotations to PubChem

PubChem contains several glycans sourced from different databases such as GlyTouCan, ChEBI, etc. Utilizing the mapping between GlyTouCan and PubChem databases, GlyGen submitted important glycan annotations such as glycan motif (e.g. Lewis X, Type 2 LN), composition (Hex5 HexNAc4 dHex1), per-methylated mass (e.g. 2221.13523366), classification (e.g. N-glycan (complex)) for glycans common between GlyGen and PubChem Compound databases (e.g. https://pubchem.ncbi.nlm.nih.gov/compound/25098607#section=Biologic-Description) Additionally, the individual monosaccharides in the composition section are further linked to the corresponding PubChem Compound pages (e.g. Hex or Hexose is linked to https://pubchem.ncbi.nlm.nih.gov/compound/206). Glycosylation annotations from GlyGen are also submitted to the “Glycobiology” section of the PubChem protein pages. Annotations such as the glycosylation site, glycosylation type (e.g. N-linked), GlyTouCan accession, glycan images (SNFG format), literature evidence in form a table is provided for associated protein entries. Additionally, a glycosylation summary stating the total number of glycosylation sites and the total number of N-liked and O-linked sites is provided at the top. E.g. https://pubchem.ncbi.nlm.nih.gov/protein/P00533#section=Glycosylation. The table displays only those glycosylation sites where the corresponding glycan is also present in the PubChem Compound database (i.e. has CID). This connectivity promotes interoperability within PubChem Protein and Compound pages. All annotations submitted by GlyGen can be found on PubChem’s data sources page (https://pubchem.ncbi.nlm.nih.gov/source/23201) or can be found at GlyGen DATA (glycan list: https://data.glygen.org/GLY_000281, glycan classification: https://data.glygen.org/GLY_000282, glycan motif: https://data.glygen.org/GLY_000283, glycan monosaccharide composition: https://data.glygen.org/GLY_000286, glycosylation annotations: https://data.glygen.org/GLY_000499).

This process enriches the existing glycan and protein data available in PubChem and connects them together through glycosylation information within the same database. Annotations submitted by GlyGen are systematically added and regularly updated to ensure synchronization between the two databases. Additionally, all the submitted annotations are bi-directionally linked at a glycan and protein level with GlyGen.

### Connecting glycans to reaction and pathway resources

Major glycoprotein databases (such as UniCarbKB, GlyConnect) use GlyTouCan accessions as primary identifiers to report glycans, whereas, reaction or pathway databases (like Rhea or Reactome) uses ChEBI Ids (See Figure 1). GlyGen utilized the GlyTouCan accession to ChEBI ID mapping to connect glycans with glycosylation annotations as well as enzymes and pathways information under one domain. E.g. in GlyGen glycan G96881BQ is connected to glycosylation information at section: https://glygen.org/glycan/G96881BQ#Associated-Protein (where information about the protein, site, source (database, literature evidence) is provided) and to reaction and pathway information through cross-references with Rhea (https://www.rhea-db.org/rhea/25580) and Reactome (https://reactome.org/content/detail/R-ALL-6787679) at section: https://glygen.org/glycan/G96881BQ#Cross-References. The Rhea cross-reference provides information on reaction, participants and literature evidence, whereas, the Reactome cross-reference provides pathways and reaction information for the compound in different species.

**Figure 1:**
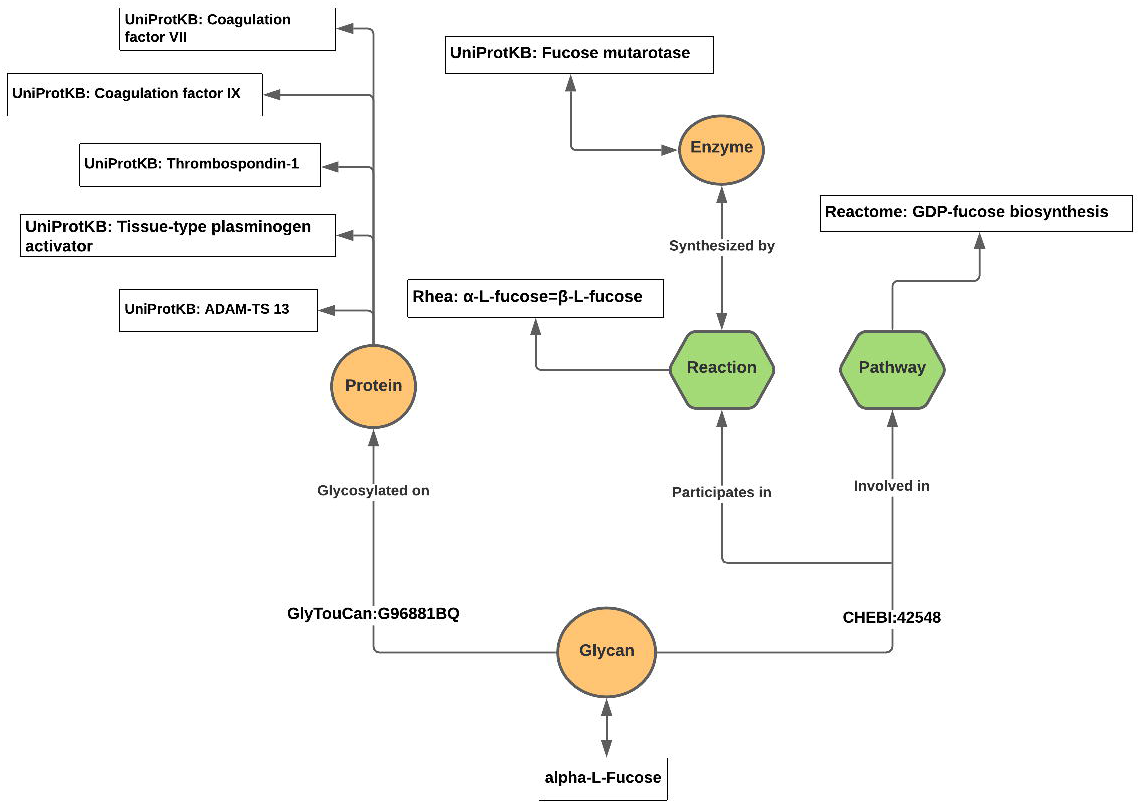
Network of the glycan annotations sourced from various databases which use either ChEBI ID or GlyTouCan accessions as primary identifiers. Utilizing the ChEBI ID to GlyTouCan accession mapping, GlyGen is able to map across multiple databases connecting glycan with information on glycosylation, reaction, and pathway (e.g. https://glygen.org/glycan/G96881BQ#Cross-References). This network is restricted to human proteins.

## DISCUSSION

Just as the usefulness of genomic and proteomic resources grew as their coverage of genes and proteins expanded, so too will the usefulness of glycan and glycosylation databases increase as the number of well-mapped glycan structures approaches comprehensive coverage. For glycans, reaching this goal has added difficulty due to the chemical nature of the compounds and the inherent limitations of the approaches for their structural analysis. Therefore, well developed submission pipelines that capture and cross-link as much information as possible are essential for growth in the glycosciences and for connecting glycan data to broader bioinformatics domains.

### Novelty

GlyTouCan, PubChem and ChEBI are major bioinformatics resources that are routinely used by many different research groups. However, the ChEBI and PubChem databases have significantly less glycan data compared to GlyTouCan. This novel pipeline allows integration of valuable glycan and glycan-related annotations hosted by GlyGen into PubChem and ChEBI databases while maintaining the links to GlyTouCan database. Additionally, ChEBI’s routine submissions to PubChem allows the newly integrated glycans to be further integrated into the PubChem database. As a result, this pipeline is not only able to identify and increase the glycan data coverage in ChEBI and PubChem databases but is also able to synchronize and map these entries ensuring connectivity. Moreover, the established connection allows further mapping and integration of other important resources (such as Rhea or Reactome) to connect reaction and pathway annotations which were previously isolated from glycoprotein data. Through this collaboration, interoperability of glycan data is achieved systematically between major databases while providing access to valuable annotations and features unique to each database.

### Applications

Currently, the glycosylation data available in the UniProt database describes the protein, glycosylation site, associated literature (for reported glycosylation) and in some cases additional description (such as high mannose, etc.) about the glycan or glycosylation (e.g. https://www.uniprot.org/uniprot/P00533#ptm_processing). However it lacks the information on the specific glycan that may be reported on a particular site on the protein. Through GlyGen’s collaboration, specific glycans (sourced from multiple resources) will be annotated to the protein and site data already present in the UniProt database (see Supplementary Figure S2). These glycans originally identified by GlyTouCan accessions by source databases and later mapped to ChEBI ID’s by GlyGen will be added to the glycoprotein sites at the ProtVista browser in UniProt. ProtVista is a protein browser providing a compact representation and access to the functional amino acid residue annotation in UniProt such as domains, sites, post-translational modifications and variants. Protvista is available through the UniProt web site and the source code is publicly available through as a web component (https://github.com/ebi-webcomponents/protvista-uniprot). The submitted glycans (identified by a ChEBI ID) along with a 2-D image will be cross-referenced to GlyGen protein pages. Additionally, UniProt users will be able to query for glycans connecting them to knowledge on proteins e.g. proteins that bind them and enzymes that synthesize or degrade them.

### Future plans

In order to sustain the growing GlyGen glycan collection, one of the major goals is to automate ~90% of the submission pipeline and reduce the time required to manually register ambiguously defined glycans. We also plan to document this pipeline so that resources like GlyTouCan, GlyConnect, and others can utilize this pipeline to integrate their glycans into the ChEBI and the PubChem databases directly or via GlyGen. Additional plans include the generation of tutorials so that users can directly register or map ambiguously-defined glycans into the ChEBI database. This can be achieved by using the PubMed ID or other publication or database identifier as a ChEBI name for the glycan entries in order to comply with the ChEBI registration protocols. Additionally, as GNOme also provides glycan-glycan relationships (using GlyTouCan accession as the primary identifier), the ambiguously defined glycans can be potentially linked to possible fully-determined, structurally-related glycans within the ChEBI database.

## MATERIALS AND METHODS

The GlyTouCan database hosts over 120,000 glycans, each identified by a GlyTouCan accession. The glycans hosted by GlyTouCan are generally deposited by various groups or individual researchers and often include glycans that are not curated and/or lack important annotations such as species, expression system, etc. As a result, GlyGen has developed an inclusion criteria in an effort to generate a subset of glycans that are unique and biologically relevant. These glycans along with valuable annotations (sourced via GlyGen’s collaborators) are available under GlyGen’s domain.

### Rules to generate GlyGen glycan set

The GlyGen glycan set (v1.5.13) comprises a subset (~24%) of the total GlyTouCan glycan set and is seeded with GlyTouCan glycan accessions annotated as human (NCBI Tax ID: 9606), mouse (NCBI Tax ID: 10090), rat (NCBI Tax ID: 10116), HCV1a (NCIB TaxID: 63746), SARS CoV 2 (NCBI TaxID:2697049) by GlyTouCan, UniCarbKB, or GlyConnect glycan data-resources; glycans representing motifs from the GlycoMotif (http://glycomotif.glyomics.org) glycan motif data-resource; GlyGen synthetic glycans; and glycans observed in human contexts in the GPTwiki (http://gptwiki.glyomics.org) glycopeptide transition data-resource. The GlyGen glycan set is then extended to include any GlyTouCan glycan accession that subsumes or shares a monosaccharide composition with a seed-glycan using the GNOme glycan naming and subsumption ontology (http://obofoundry.org/ontology/gno, http://gnome.glyomics.org). (See Supplementary Figure S3). GlyGen data processing and integration framework is utilized to quality control and standardized this data which is supplied to GlyGen portal (Kahsay et al. 2020).

### Harmonization of glycans between different databases

GlyTouCan and ChEBI databases routinely submit entities (as MDL SDF files) to PubChem which are classified as “substances” and identified by unique substance or SIDs. The standardization process then validates and normalizes the chemical entities to generate a PubChem compound database where the standardized compound is identified by a unique CID (Kim et al. 2016). Multiple SIDs can map to the same CID (see Figure 2).

**Figure 2:**
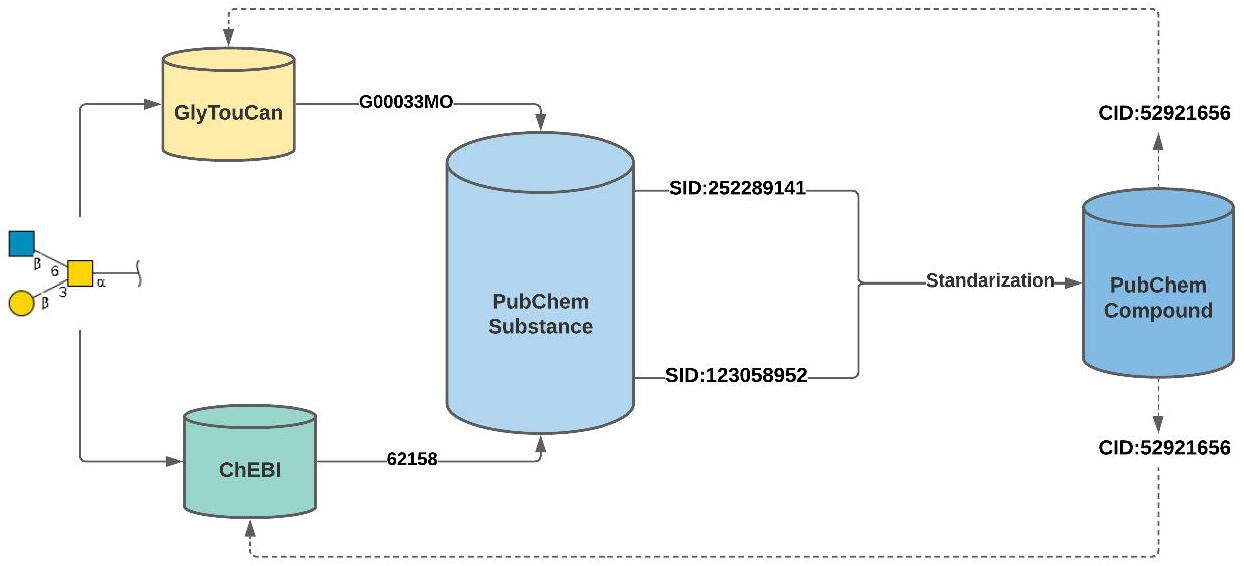
Outlines the data flow of glycans across GlyTouCan, ChEBI, and PubChem databases. The figure shows an example of the same glycan (beta-D-Galp-(1->3)-[beta-D-GlcpNAc-(1->6)]-alpha-D-GalpNAc) present in GlyTouCan (G00033MO) and ChEBI (62158) under respective database identifiers. The GlyTouCan accession and ChEBI ID is mapped to unique PubChem Substance identifiers (G00033MO to SID:252289141; CHEBI:62158 to SID:123058952) when submitted to the PubChem database. PubChem’s standardization process maps both the SID’s to a single compound identifier (CID:52921656). The same CID is utilized as a cross-reference by both GlyTouCan and ChEBI databases.

PubChem only accepts glycans where the arrangement of the monosaccharides, glycosidic bonds and linkages or anomeric positions are known. However, ambiguity regarding the first linkage (between the glycan and conjugate) and stereochemistry of any particular atom is tolerated within PubChem. As a result, not all GlyTouCan accessions are included in the PubChem database. Additionally, before the start of the collaboration only a small portion of the available glycans in PubChem were mapped to the ChEBI database resulting in low connectivity across GlyTouCan, PubChem and ChEBI databases.

As an attempt to harmonize glycans between GlyTouCan and ChEBI, the GlyGen glycan set (v.1.5.13) comprising 29,290 glycans was identified as the primary list. The set contains 19,039 glycans associated with saccharide subsumption category as defined by the Glycan Naming Ontology or GNOme (for description of GNOme classes see Supplementary Figure S1). Rest includes glycans associated with topology (6143), composition (2652), base composition (1386) and undefined (97) categories. The set also includes 13,389 fully-determined glycans that belong to either saccharide, topology or composition subsumption categories. An integration pipeline was developed to map and register all GlyGen glycans into the ChEBI database (see Figure 3).

**Figure 3:**
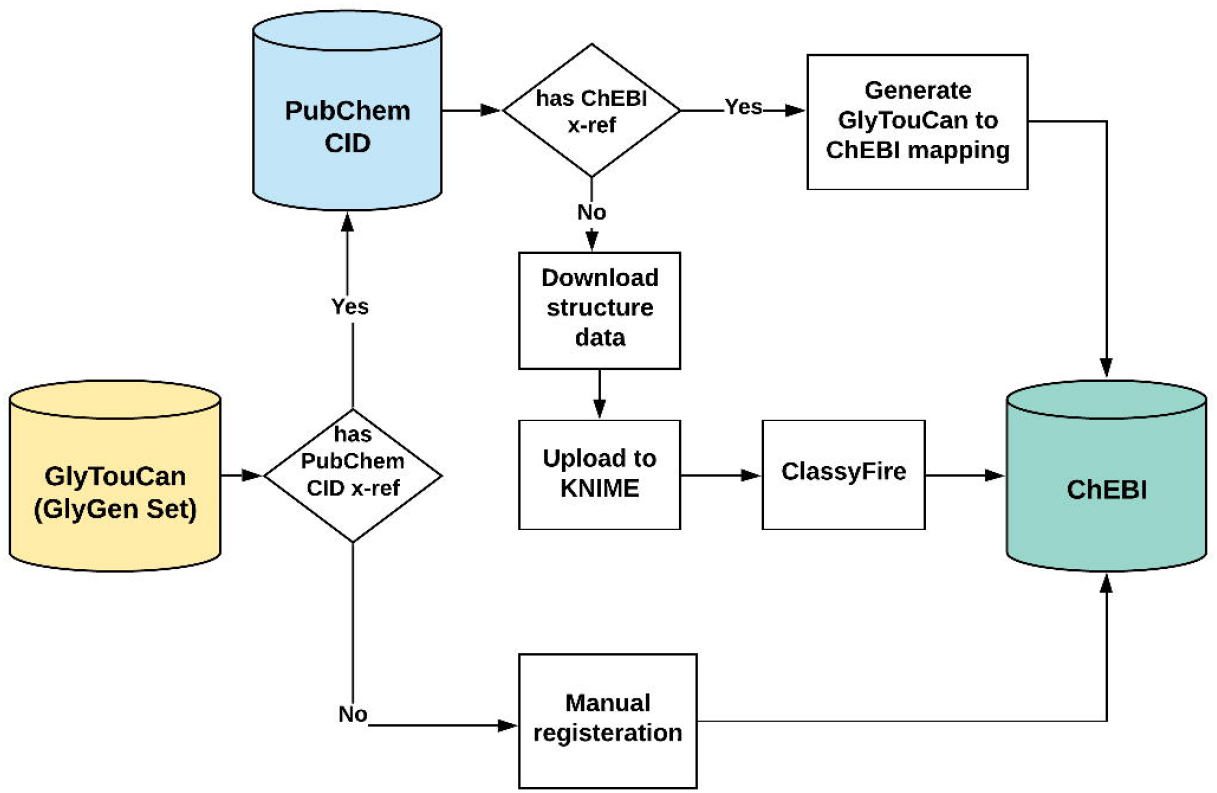
Overview of the data integration pipeline to map or register the GlyGen glycan set of 29,290 GlyTouCan accessions into the ChEBI database. If the GlyTouCan accession had a PubChem CID and a corresponding ChEBI ID, then a cross-reference mapping was generated and added to the ChEBI database where the corresponding GlyTouCan accession was added as a cross-reference. GlyTouCan accessions with a PubChem CID but without a ChEBI ID were uploaded to ChEBI using applications like KNIME and ClassyFire. The remaining GlyTouCan accessions where a PubChem CID mapping was missing were manually registered in the ChEBI database.

#### Using PubChem as an anchor database

PubChem Compound was used as an anchor database to identify the existing mapping between the GlyTouCan and ChEBI databases. Out of the GlyGen glycan list of 29,290 GlyTouCan accessions 10,496 were already cross-referenced to PubChem compound identifiers (CIDs). These mostly included glycans which are fully-determined with or without known first linkage (between the glycan and conjugate). This set also includes glycans which consists of a single monosaccharide as well glycans with one or more undefined atomic stereocenter. This mapping was utilized to retrieve the corresponding PubChem CID to ChEBI ID mapping (e.g. using the mapping between GlyTouCan G14047PA to PubChem CID 70788991 and ChEBI:72002 to PubChem CID 70788991, the GlyTouCan accession was mapped to the ChEBI ID). As a result, over 1,700 glycans were mapped between the GlyTouCan and ChEBI databases. However, this method provided only about 5% coverage for the GlyGen glycan set. As for the remaining PubChem CIDs where a ChEBI ID mapping was not directly available from the PubChem database, the integration pipeline was modified so that the structure data for individual entries (downloaded from PubChem) could be uploaded into applications like KNIME (Fillbrunn et al. 2017) and ClassyFire (Djoumbou Feunang et al. 2016) to generate a suitable SD file for ChEBI. In this process, the downloaded PubChem structure data for individual entries were uploaded to the KNIME application where the data was first cleaned (e.g. remove explicit hydrogens while maintaining the stereochemical integrity of the structure) and the additional information (such as associated names, synonyms and WURCS sequence) was extracted. The data was further uploaded to the ClassyFire application (at 12 requests per minute rate) in order to generate ontology terms. The generated ontology terms were then converted to the best matching ChEBI ontology terms and an SD file for ChEBI upload was generated. A default classification value (such as carbohydrates and carbohydrates-derivatives) was added in cases where ClassyFire was unable to generate an ontology term for a record. The files generated by the KNIME application were submitted to the ChEBI loader tool (in batches of 250 records) and the GlyTouCan accession (via PubChem CID) was either mapped to an existing ChEBI ID (if the structure was already present in ChEBI) or registered as a new entry with 2-star status. This method allowed integration of any GlyTouCan accession (within GlyGen) with a PubChem CID into the ChEBI database.

#### Manual registration of glycans into ChEBI

The PubChem Compound database facilitated mapping or registration of roughly 30% of the GlyGen glycans (GlyTouCan accessions) into the ChEBI database. The remaining set of 18,794 glycans lacked a PubChem CID, likely as they did not fit the PubChem requirements. This includes (but not restricted to) glycans that strictly compositions (e.g. G00058XR or G00057LF) or glycans with undefined stereochemistry (e.g G00318AV or G00012RZ). However, this set also includes about 3256 fully-determined glycans which lack a PubChem CID probably due to a technical error or a database synchronization issue between PubChem and the GlyTouCan database.

To establish and optimize a protocol, twenty-five high-value glycans (GlyTouCan accession) were selected to be manually registered into ChEBI (see Supplementary Table SI) as a proof of concept. The subset was identified based on the following conditions: 1) Top-ten composition glycans with the largest number of associated proteins in GlyGen (v1.5.36). 2) Top-ten ambiguously-defined glycans with the largest number of associated proteins in GlyGen (v1.5.36) and 3) Five composition glycans associated to the HCV1a protein in GlyGen (v1.5.36). Based on the following approaches the remaining glycans will be registered manually: 1) ***Fully determined glycans:*** Such glycans can be treated as ordinary chemical entities in the same manner as most of the other entities in the ChEBI database. Entries can be added by a ChEBI curator using the Curator tool, or by individual submitters using the ChEBI submissions tool (in which a submitter provides a name, structure, etc.). Each submission is immediately visible on the public website as a 2-star (unchecked) entry and is upgraded to 3-star (checked) status after being checked by a ChEBI curator. After being indexed overnight, submissions can be fully searched, for example, by name, synonyms, InChI, InChIKey, SMILES, and structure, the following day. 2) ***Glycans with unknown stereochemistry:*** In principle, all of the methods described for fully determined structures can be used in cases where some (or all!) of the stereochemistry is unknown. If the alpha/beta configuration of one or more glycosidic bonds in a glycan is not known, then such bonds are depicted in the chemical structure as normal single bonds. For double bonds of unknown configuration, ‘wavy’ bonds can be used to link the double bond to one or more of its substituents. 3) ***Glycans with unknown position of attachment:*** If the position of attachment of a glycosyl group to a glycan is not known, then the structure is drawn using “R” atoms to indicate the possible sites of attachment. In the Definition field, a note is included to indicate the possible values of R at each position 4) ***Compositions:*** In cases where only the composition such as the type (hexose, pentose, etc.) and number of monosaccharides is known then such glycans can be registered using the source (in this case GlyTouCan) accession as a ChEBI name and the composition of a glycan (e.g. Composition: Hex2 HexNAc5.) can be added as part of the description. However, due to high number of possible linkages, such entries would lack a ChEBI image.

#### Integration of glycan annotations to PubChem

A sustainable approach was developed to submit glycan and glycoprotein annotations from GlyGen to PubChem Compound and protein pages. After registering GlyGen as a user through PubChem upload mechanism (https://pubchem.ncbi.nlm.nih.gov/upload/) stable unique URLs of annotation specific GlyGen datasets (e.g. https://data.glygen.org/GLY_000283) are regularly and automatically consumed by the PubChem database to add and update the annotations. Each compound in PubChem has a dedicated page and is identified with a unique compound identifier (or CID) which represents a standardized entry from multiple substances submitted to PubChem. Utilizing the existing GlyTouCan accession to PubChem CID mapping and GlyGen’s annotation specific dataset, annotations such as GlyGen glycan list (https://data.glygen.org/GLY_000281), glycan classification (https://data.glygen.org/GLY_000282), glycan motif (https://data.glygen.org/GLY_000283), and monosaccharide composition (https://data.glygen.org/GLY_000286) were added to the respective PubChem Compound page. The annotations were added under the ‘Biologic Description’ section on the PubChem Compound page along with the URL for the GlyGen glycan page. The individual monosaccharides within the glycan composition are further linked to respective entries within PubChem Compound pages. Similar to the Compound pages, PubChem has dedicated pages for the proteins identified by the NCBI Protein accessions. Utilizing the UniProtKB accession, GlyGen provided the respective glycoprotein annotations (using dataset https://data.glygen.org/GLY_000499) to the PubChem Protein pages. The annotations are added under the ‘Glycobiology’ section along with a link to GlyGen Protein pages. The section also contains a table that shows glycosylation information including glycosylation site, glycosylation type, GlyTouCan ID, glycan image, PubChem CID, and PMID evidence for only a subset of glycoprotein annotations where the associated glycan (GlyTouCan accession) has a corresponding PubChem CID for a given glycoprotein.

## ACKNOWLEDGEMENTS

Kiyoko Aoki-Kinoshita, Terence Murphy and Andrew Leach

## ABBREVIATIONS

EMBL-EBI: EMBL-European Bioinformatics Institute
SIB: Swiss Institute of Bioinformatics
ChEBI: Chemical Entities of Biological Interest
CFG: Consortium for Functional Glycomics
GNOme: Glycan Naming Ontology
CID: PubChem Compound Identifier
SID: PubChem Substance Identifier
WURCS: Web3 Unique Representation of Carbohydrate Structures
SMILES: Simplified Molecular-Input Line-Entry System
InChl: International Chemical Identifier
IUPAC: International Union of Pure and Applied Chemistry
HCV: Hepatitis C virus
X-ref: Cross-reference

## DATA AVAILABILITY STATEMENT

The data can be openly accessed through the GlyGen database at https://www.glygen.org/ and https://data.glygen.org/

## REFERENCES

Acevedo A, Simister R, McQueen-Mason SJ, Gomez LD. 2019. Sudangrass, an alternative lignocellulosic feedstock for bioenergy in Argentina. PLoS One 14(5):e0217435

Alocci D, Mariethoz J, Gastaldello A, Gasteiger E, Karlsson NG, Kolarich D, Packer NH, Lisacek F. 2019. GlyConnect: Glycoproteomics Goes Visual, Interactive, and Analytical. J Proteome Res 18(2):664–677

Ashburner M, Ball CA, Blake JA, Botstein D, Butler H, Cherry JM, Davis AP, Dolinski K, Dwight SS, Eppig JT et al. . 2000. Gene ontology: tool for the unification of biology. The Gene Ontology Consortium. Nat Genet 25(1):25–29

Bohm M, Bohne-Lang A, Frank M, Loss A, Rojas-Macias MA, Lutteke T. 2019. Glycosciences.DB: an annotated data collection linking glycomics and proteomics data (2018 update). Nucleic Acids Res 47(D1):D1195–D1201

Bonifacino S, Resquin F, Lopretti M, Buxedas L, Vazquez S, Gonzalez M, Sapolinski A, Hirigoyen A, Doldan J, Rachid C et al. . 2021. Bioethanol production using high density Eucalyptus crops in Uruguay. Heliyon 7(1):e06031

Campbell MP, Peterson R, Mariethoz J, Gasteiger E, Akune Y, Aoki-Kinoshita KF, Lisacek F, Packer NH. 2014. UniCarbKB: building a knowledge platform for glycoproteomics. Nucleic Acids Res 42(Database issue):D215–221

Chen K, Gentry-Maharaj A, Burnell M, Steentoft C, Marcos-Silva L, Mandel U, Jacobs I, Dawnay A, Menon U, Blixt O. 2013. Microarray Glycoprofiling of CA125 improves differential diagnosis of ovarian cancer. J Proteome Res 12(3):1408–1418

Clerc O, Deniaud M, Vallet SD, Naba A, Rivet A, Perez S, Thierry-Mieg N, Ricard-Blum S. 2019. MatrixDB: integration of new data with a focus on glycosaminoglycan interactions. Nucleic Acids Res 47(D1):D376–D381

Clifton-Brown J, Harfouche A, Casler MD, Dylan Jones H, Macalpine WJ, Murphy-Bokern D, Smart LB, Adler A, Ashman C, Awty-Carroll D et al. . 2019. Breeding progress and preparedness for mass-scale deployment of perennial lignocellulosic biomass crops switchgrass, miscanthus, willow and poplar. Glob Change Biol Bioenergy 11(1):118–151

Copeland R, Balasubramaniam A, Tiwari V, Zhang F, Bridges A, Linhardt RJ, Shukla D, Liu J. 2008. Using a 3-O-sulfated heparin octasaccharide to inhibit the entry of herpes simplex virus type 1. Biochemistry 47(21):5774–5783

Council NR. 2012. Transforming Glycoscience: A Roadmap for the Future. Transforming Glycoscience: A Roadmap for the Future. Washington (DC).

Crabtree MB, Kinney RM, Miller BR. 2005. Deglycosylation of the NS1 protein of dengue 2 virus, strain 16681: construction and characterization of mutant viruses. Arch Virol 150(4):771–786

Crispin M, Ward AB, Wilson IA. 2018. Structure and Immune Recognition of the HIV Glycan Shield. Annu Rev Biophys 47:499–523

Cummings RD, Pierce JM. 2014. The challenge and promise of glycomics. Chem Biol 21(1):1–15

da Costa RM, Pattathil S, Avci U, Lee SJ, Hazen SP, Winters A, Hahn MG, Bosch M. 2017. A cell wall reference profile for Miscanthus bioenergy crops highlights compositional and structural variations associated with development and organ origin. New Phytol 213(4):1710–1725

Daus S, Heinze T. 2010. Xylan-based nanoparticles: prodrugs for ibuprofen release. Macromol Biosci 10(2):211–220

Degtyarenko K, de Matos P, Ennis M, Hastings J, Zbinden M, McNaught A, Alcantara R, Darsow M, Guedj M, Ashburner M. 2008. ChEBI: a database and ontology for chemical entities of biological interest. Nucleic Acids Res 36(Database issue):D344–350

Djoumbou Feunang Y, Eisner R, Knox C, Chepelev L, Hastings J, Owen G, Fahy E, Steinbeck C, Subramanian S, Bolton E et al. . 2016. ClassyFire: automated chemical classification with a comprehensive, computable taxonomy. J Cheminform 8:61

Figueiredo P, Lintinen K, Kiriazis A, Hynninen V, Liu Z, Bauleth-Ramos T, Rahikkala A, Correia A, Kohout T, Sarmento B et al. . 2017. In vitro evaluation of biodegradable lignin-based nanoparticles for drug delivery and enhanced antiproliferation effect in cancer cells. Biomaterials 121:97–108

Fillbrunn A, Dietz C, Pfeuffer J, Rahn R, Landrum GA, Berthold MR. 2017. KNIME for reproducible cross-domain analysis of life science data. J Biotechnol 261:149–156

Gene Ontology Consortium. 2021. The Gene Ontology resource: enriching a GOld mine. Nucleic Acids Res 49(D1):D325–D334

Hansen NM, Plackett D. 2008. Sustainable films and coatings from hemicelluloses: a review. Biomacromolecules 9(6):1493–1505

Hartman J, Albertsson A-C, Lindblad MS, Sjöberg J. 2006. Oxygen barrier materials from renewable sources: Material properties of softwood hemicellulose-based films. Journal of Applied Polymer Science 100(4):2985–2991

Hashimoto K, Goto S, Kawano S, Aoki-Kinoshita KF, Ueda N, Hamajima M, Kawasaki T, Kanehisa M. 2006. KEGG as a glycome informatics resource. Glycobiology 16(5):63R–70R

Herget S, Ranzinger R, Maass K, Lieth CW. 2008. GlycoCT-a unifying sequence format for carbohydrates. Carbohydr Res 343(12):2162–2171

Jassal B, Matthews L, Viteri G, Gong C, Lorente P, Fabregat A, Sidiropoulos K, Cook J, Gillespie M, Haw R et al. . 2020. The reactome pathway knowledgebase. Nucleic Acids Res 48(D1):D498–D503

Kahsay R, Vora J, Navelkar R, Mousavi R, Fochtman BC, Holmes X, Pattabiraman N, Ranzinger R, Mahadik R, Williamson T et al. . 2020. GlyGen data model and processing workflow. Bioinformatics 36(12):3941–3943

Kim S, Chen J, Cheng T, Gindulyte A, He J, He S, Li Q, Shoemaker BA, Thiessen PA, Yu B et al. . 2021. PubChem in 2021: new data content and improved web interfaces. Nucleic Acids Res 49(D1):D1388–D1395

Kim S, Thiessen PA, Bolton EE, Chen J, Fu G, Gindulyte A, Han L, He J, He S, Shoemaker BA et al. . 2016. PubChem Substance and Compound databases. Nucleic Acids Res 44(D1):D1202–1213

Kirwan A, Utratna M, O’Dwyer ME, Joshi L, Kilcoyne M. 2015. Glycosylation-Based Serum Biomarkers for Cancer Diagnostics and Prognostics. Biomed Res Int 2015:490531

Lemus R, Parrish DJ, Wolf DD. 2014. Switchgrass cultivar/ecotype selection and management for biofuels in the upper southeast USA. ScientificWorldJournal 2014:937594

Li W, Hulswit RJG, Widjaja I, Raj VS, McBride R, Peng W, Widagdo W, Tortorici MA, van Dieren B, Lang Y et al. . 2017. Identification of sialic acid-binding function for the Middle East respiratory syndrome coronavirus spike glycoprotein. Proc Natl Acad Sci U S A 114(40):E8508–E8517

Lin B, Qing X, Liao J, Zhuo K. 2020. Role of Protein Glycosylation in Host-Pathogen Interaction. Cells 9(4)

Lombardot T, Morgat A, Axelsen KB, Aimo L, Hyka-Nouspikel N, Niknejad A, Ignatchenko A, Xenarios I, Coudert E, Redaschi N et al. . 2019. Updates in Rhea: SPARQLing biochemical reaction data. Nucleic Acids Res 47(D1):D596–D600

Matsubara M, Aoki-Kinoshita KF, Aoki NP, Yamada I, Narimatsu H. 2017. WURCS 2.0 Update To Encapsulate Ambiguous Carbohydrate Structures. J Chem Inf Model 57(4):632–637

Miller JL, de Wet BJ, Martinez-Pomares L, Radcliffe CM, Dwek RA, Rudd PM, Gordon S. 2008. The mannose receptor mediates dengue virus infection of macrophages. PLoS Pathog 4(2):e17

Neelamegham S, Aoki-Kinoshita K, Bolton E, Frank M, Lisacek F, Lutteke T, O’Boyle N, Packer NH, Stanley P, Toukach P et al. . 2019. Updates to the Symbol Nomenclature for Glycans guidelines. Glycobiology 29(9):620–624

O’Neill MA, Moon RJ, York WS, Darvill AG. 2015. Glycans in Bioenergy and Materials Science. In: rd, Varki A, Cummings RD, Esko JD, Stanley P, Hart GW, Aebi M, Darvill AG, Kinoshita T, Packer NH et al., editors. Essentials of Glycobiology. Cold Spring Harbor (NY). p. 755–760.

Panico M, Bouche L, Binet D, O’Connor MJ, Rahman D, Pang PC, Canis K, North SJ, Desrosiers RC, Chertova E et al. . 2016. Mapping the complete glycoproteome of virion-derived HIV-1 gp120 provides insights into broadly neutralizing antibody binding. Sci Rep 6:32956

Patel JA, Dean DA, King CH, Xiao N, Koc S, Minina E, Golikov A, Brooks P, Kahsay R, Navelkar R et al. . 2021. Bioinformatics tools developed to support BioCompute Objects. Database (Oxford) 2021

Penades S, Davis BG, Seeberger PH. 2015. Glycans in Nanotechnology. In: rd, Varki A, Cummings RD, Esko JD, Stanley P, Hart GW, Aebi M, Darvill AG, Kinoshita T, Packer NH et al., editors. Essentials of Glycobiology. Cold Spring Harbor (NY). p. 743–753.

Reily C, Stewart TJ, Renfrow MB, Novak J. 2019. Glycosylation in health and disease. Nat Rev Nephrol 15(6):346–366

Sai Prasanna N, Mitra J. 2020. Isolation and characterization of cellulose nanocrystals from Cucumis sativus peels. Carbohydr Polym 247:116706

Schmer MR, Vogel KP, Mitchell RB, Perrin RK. 2008. Net energy of cellulosic ethanol from switchgrass. Proc Natl Acad Sci U S A 105(2):464–469

Seeberger PH, Cummings RD. 2015. Glycans in Biotechnology and the Pharmaceutical Industry. In: rd, Varki A, Cummings RD, Esko JD, Stanley P, Hart GW, Aebi M, Darvill AG, Kinoshita T, Packer NH et al., editors. Essentials of Glycobiology. Cold Spring Harbor (NY). p. 729–741.

Sharthiya H, Seng C, Van Kuppevelt TH, Tiwari V, Fornaro M. 2017. HSV-1 interaction to 3-O-sulfated heparan sulfate in mouse-derived DRG explant and profiles of inflammatory markers during virus infection. J Neurovirol 23(3):483–491

Shiraki K, Takase K, Tameda Y, Hamada M, Kosaka Y, Nakano T. 1995. A clinical study of lectin-reactive alpha-fetoprotein as an early indicator of hepatocellular carcinoma in the follow-up of cirrhotic patients. Hepatology 22(3):802–807

Simonyan V, Goecks J, Mazumder R. 2017. Biocompute Objects-A Step towards Evaluation and Validation of Biomedical Scientific Computations. PDA J Pharm Sci Technol 71(2):136–146

The UniProt Consortium. 2021. UniProt: the universal protein knowledgebase in 2021. Nucleic Acids Res 49(D1):D480–D489

Tiemeyer M, Aoki K, Paulson J, Cummings RD, York WS, Karlsson NG, Lisacek F, Packer NH, Campbell MP, Aoki NP et al. . 2017. GlyTouCan: an accessible glycan structure repository. Glycobiology 27(10):915–919

Varki A. 2017. Biological roles of glycans. Glycobiology 27(1):3–49

Varki A, Cummings RD, Aebi M, Packer NH, Seeberger PH, Esko JD, Stanley P, Hart G, Darvill A, Kinoshita T et al. . 2015. Symbol Nomenclature for Graphical Representations of Glycans. Glycobiology 25(12):1323–1324

Varki A, Gagneux P. 2015. Biological Functions of Glycans. In: rd, Varki A, Cummings RD, Esko JD, Stanley P, Hart GW, Aebi M, Darvill AG, Kinoshita T, Packer NH et al., editors. Essentials of Glycobiology. Cold Spring Harbor (NY). p. 77–88.

Walls AC, Tortorici MA, Frenz B, Snijder J, Li W, Rey FA, DiMaio F, Bosch BJ, Veesler D. 2016. Glycan shield and epitope masking of a coronavirus spike protein observed by cryoelectron microscopy. Nat Struct Mol Biol 23(10):899–905

Wang W, Owen SM, Rudolph DL, Cole AM, Hong T, Waring AJ, Lal RB, Lehrer RI. 2004. Activity of alpha- and theta-defensins against primary isolates of HIV-1. J Immunol 173(1):515–520

Watanabe Y, Allen JD, Wrapp D, McLellan JS, Crispin M. 2020. Site-specific glycan analysis of the SARS-CoV-2 spike. Science 369(6501):330–333

Wijaya CJ, Ismadji S, Gunawan S. 2021. A Review of Lignocellulosic-Derived Nanoparticles for Drug Delivery Applications: Lignin Nanoparticles, Xylan Nanoparticles, and Cellulose Nanocrystals. Molecules 26(3)

York WS, Mazumder R, Ranzinger R, Edwards N, Kahsay R, Aoki-Kinoshita KF, Campbell MP, Cummings RD, Feizi T, Martin M et al. . 2020. GlyGen: Computational and Informatics Resources for Glycoscience. Glycobiology 30(2):72–73

