## Supplementary Figure S1 for "Enhancing the interoperability of glycan data flow between ChEBI, PubChem, and GlyGen"

**Supp. Fig. 1**

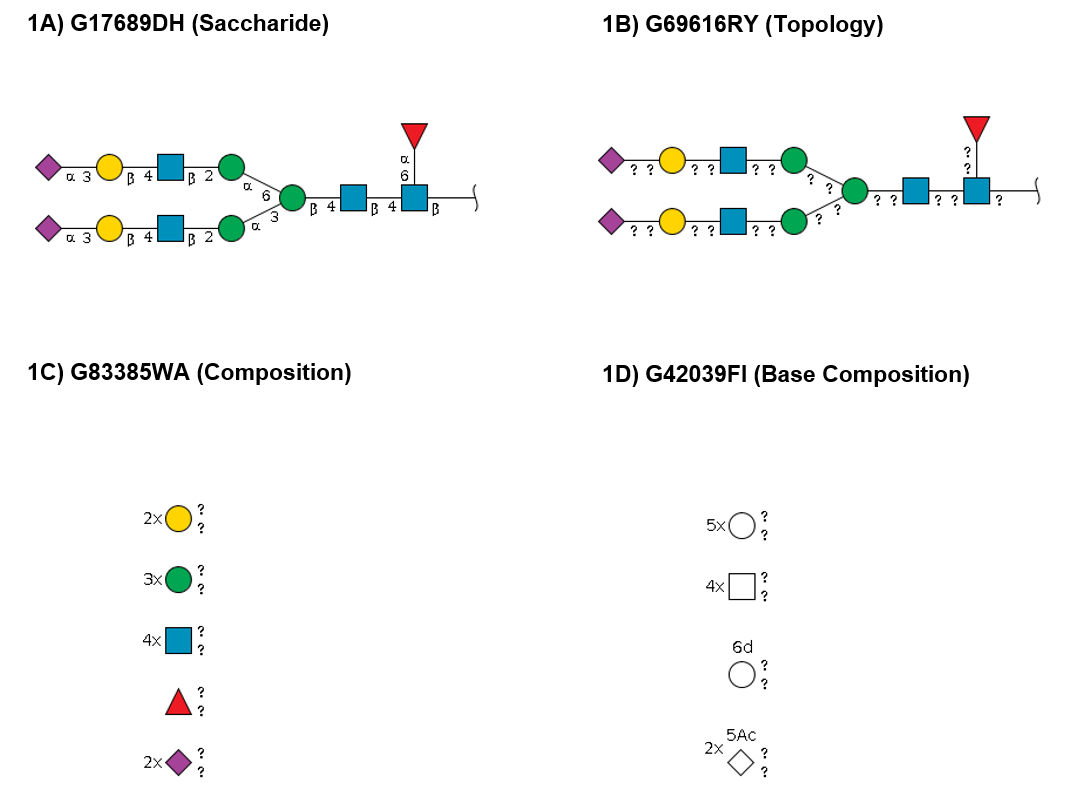

**Figure S1.** Different types of glycans (identified by a GlyTouCan accession) represented in an SNFG format with varying levels of knowledge about its stereochemistry, glycosidic bonds, anomeric configuration, etc. The types of glycan shown in the figure also describe different instances or subsumption categories in the Glycan Naming Ontology (<http://www.obofoundry.org/ontology/gno.html>) or GNOme. 1a) Shows a type of glycan (G17689DH) where the arrangement (type, number, and stereochemistry) of monosaccharides is known along with the glyosidic bonds with partial or complete information about the linkage or anomeric configuration. Such glycans are categorized under “Saccharide” (<http://purl.obolibrary.org/obo/GNO_00000016>) instance in GNOme. 1b) Shows a type of glycan (G69616RY) where there is no information about the linkage or anomeric configuration, but the arrangement of monosaccharides and glycosidic bonds is known. Such glycans belong to the “Topology” (<http://purl.obolibrary.org/obo/GNO_00000015>) instance in GNOme. 1c) Shows a type of glycan (G83385WA) where only the information about type and number of monosaccharides is known in addition to complete or partial knowledge of the stereochemistry of the involved monosaccharides. Such glycans belong to the “composition” (<http://purl.obolibrary.org/obo/GNO_00000014>) instance in GNOme. Certain glycans from these three instances are also identified as fully-determined glycans where there is complete knowledge of the arrangement of the monosaccharides, glycosidic bonds, and linkage/anomeric configuration with the exception of first linkage (between glycan and conjugate) where the ambiguity is tolerated. 1d) Shows a type of glycan (G42039FI) where only the information about the type and number of monosaccharides is known. Such glycans are categorized under the “basecomposition” (<http://purl.obolibrary.org/obo/GNO_00000013>) instance in GNOme.
