## Supplementary Figure S2 for "Enhancing the interoperability of glycan data flow between ChEBI, PubChem, and GlyGen"

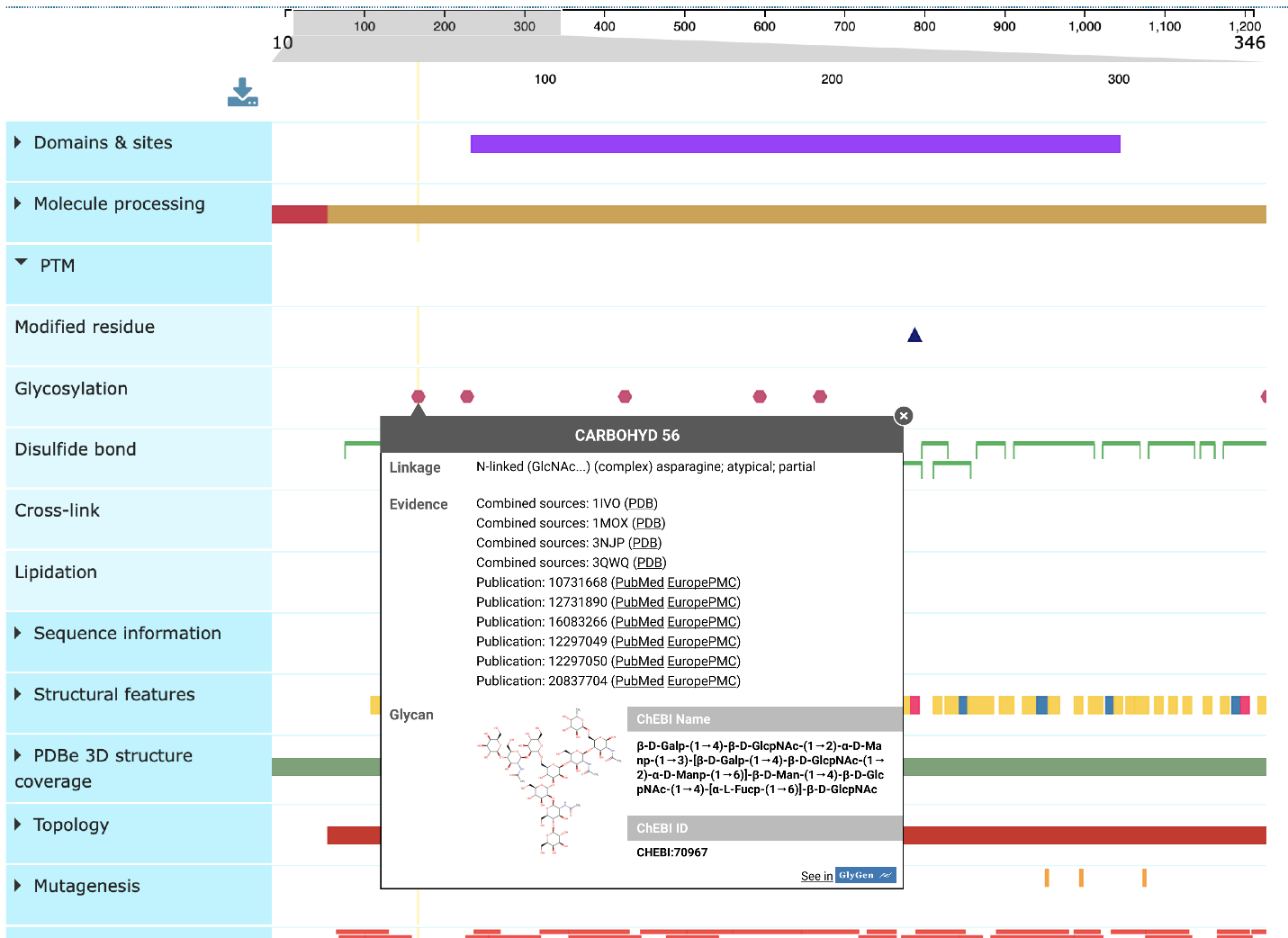

**Figure S2**. Prototype developed by GlyGen and UniProtKB to display glycan information in the UniProt’s ProtVista protein viewer. Glycans annotated in ChEBI by GlyGen will be displayed in the UniProt resource through the ProtVista viewer. This example shows the glycosylation site (at amino acid 56) of the human epidermal growth factor receptor (EGFR, UniProt accession P00533). Glycosylation sites are annotated in UniProt with the type of linkage which binds to the carbohydrate chain i.e N-linked and the reducing sugar. The evidence for the annotation i.e. literature references are also provided. The glycan annotation to the corresponding ChEBI identifier and structure will be imported from the GlyGen and ChEBI.
