## Supplementary Figure S3 for "Enhancing the interoperability of glycan data flow between ChEBI, PubChem, and GlyGen"

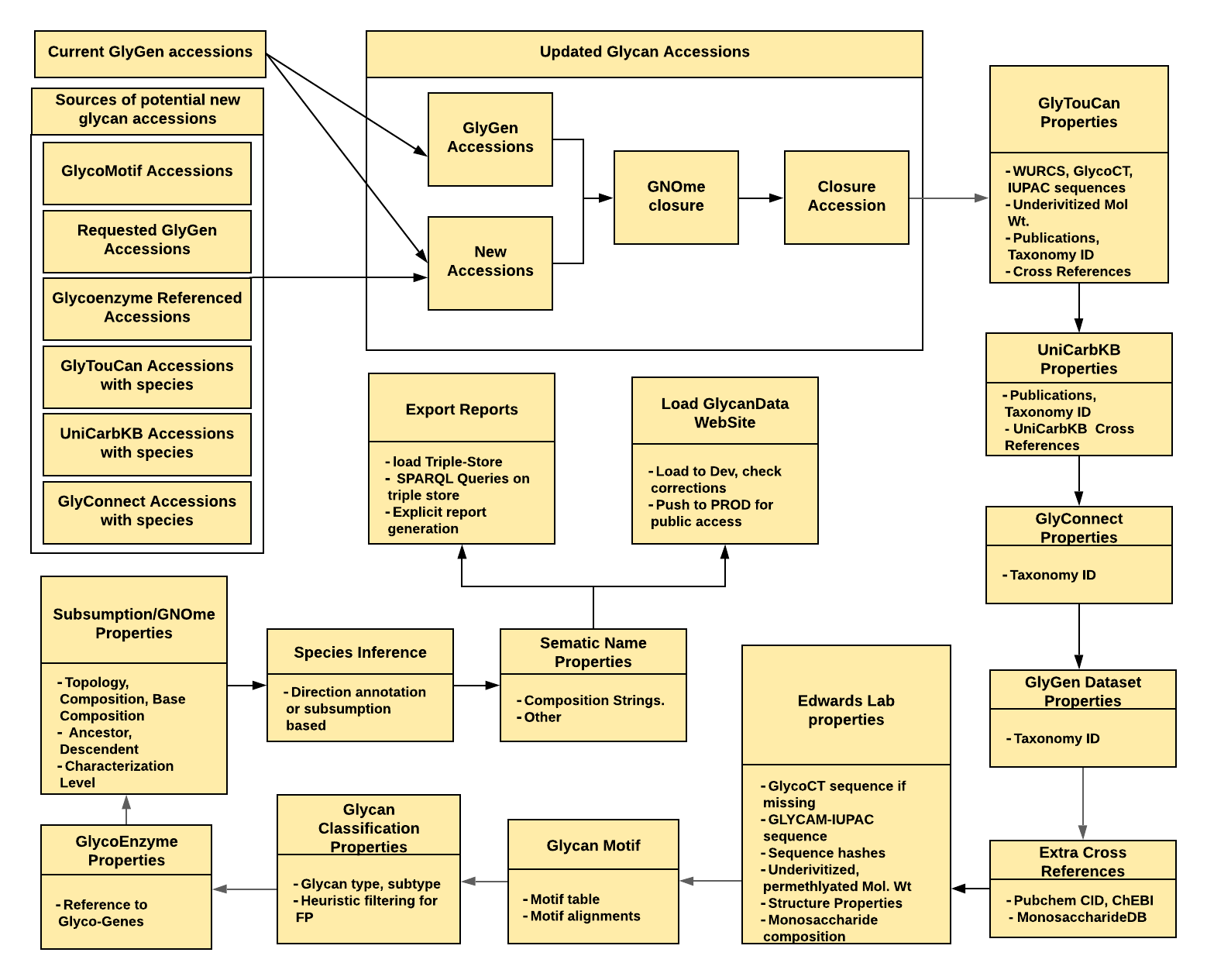

**Figure S3.** Glycan data integration from multiple resources to generate a GlyGen glycan set. Provides an overview of the steps and workflow on collection and integration of glycans from multiple glycans and glycoprotein resources as well as Glycan Naming Ontology to generate a comprehensive glycan dataset for GlyGen.
