## Supplementary Table S1 for "Enhancing the interoperability of glycan data flow between ChEBI, PubChem, and GlyGen"

**Supplementary Table 1:** Depicts the high-value, ambiguously defined glycans manually registered into the ChEBI database as a proof of concept.

| GlyTouCan Accession | Subsumption category (GNOme) | CHEBI ID | No of associated glycoproteins |
| --- | --- | --- | --- |
| G06110VR | Base Composition | CHEBI:156559 | 762 |
| G43417UB | Base Composition | CHEBI:156560 | 608 |
| G70101JE | Base Composition | CHEBI:156561 | 489 |
| G84452RH | Composition | CHEBI:156562 | 360 |
| G06356OH | Composition | CHEBI:156563 | 280 |
| G74724QE | Base Composition | CHEBI:156564 | 221 |
| G29068FM | Base Composition | CHEBI:156565 | 215 |
| G23863VK | Base Composition | CHEBI:156566 | 200 |
| G23294PN | Composition | CHEBI:156568 | 190 |
| G14669DU | Base Composition | CHEBI:156569 | 190 |
| G72291OX | Composition | CHEBI:156570 | 100 |
| G14548ZL | Saccharide | CHEBI:156581 | 40 |
| G42358LZ | Saccharide | CHEBI:156582 | 35 |
| G22140GZ | Saccharide | CHEBI:156583 | 30 |
| G40608DF | Saccharide | CHEBI:156584 | 19 |
| G70994MS | Topology | CHEBI:18133 | 17 |
| G57818FI | Saccharide | CHEBI:156585 | 13 |
| G39397SW | Saccharide | CHEBI:157599 | 12 |
| G93860XO | Saccharide | CHEBI:157638 | 10 |
| G48975GN | Saccharide | CHEBI:157639 | 10 |
| G62765YT | Base Composition | CHEBI:157633 | 9 |
| G80920RR | Base Composition | CHEBI:157634 | 9 |
| G31852PQ | Base Composition | CHEBI:157635 | 8 |
| G41247ZX | Base Composition | CHEBI:157636 | 8 |
| G02815KT | Base Composition | CHEBI:157637 | 7 |
